# Ser14-phosphorylated Rpn6 Limits Proteostasis Impairment and Pathology in Both Brain and Heart of Tauopathy Mice

**DOI:** 10.1101/2025.03.24.645024

**Authors:** Saima Ejaz, Jack O. Sternburg, Khosrow Rezvani, Md. Salim Ahammed, Samiksha Giri, Jinbao Liu, Hongmin Wang, Xuejun Wang

## Abstract

Alzheimer’s disease (AD) patients often display neurobehavioral and cardiac impairments, but the underlying factors remain unclear. Ser14 phosphorylation in RPN6 (p-S14-RPN6) mediates the activation of 26S proteasomes by protein kinase A (PKA). Proteasome priming is implicated in protection by cAMP-PKA against AD, but this remains to be established. Hence, this study was conducted to interrogate homeostatic p-S14-RPN6 in AD. The recently validated Rpn6^S14A^ knock-in (S14A) mice were crossbred with the PS19 tauopathy mice (RRID: IMSR_JAX:008169). The resultant wild type (WT), PS19, and PS19::S14A littermates were compared. Expedited declines in cognitive and motor functions as indicated respectively by significant decreases in object recognition and discrimination indexes and rotarod time were observed in PS19::S14A mice vs. PS19 mice, which is associated with more pronounced synaptic losses, microglial activation, and gliosis in the hippocampus. Compared with WT and PS19 mice, PS19::S14A mice showed exacerbated cardiac malfunction, cardiac hypertrophic responses and fibrosis, and greater increases of total and hyperphosphorylated tau proteins and ubiquitin conjugates in both hippocampi and hearts. These findings demonstrate that genetic blockade of p-S14-RPN6 exacerbates tauopathy in both the brain and heart, which for the first time establishes that homeostatic p-S14-RPN6 promotes proteostasis and protects against pathogenesis in AD.

## INTRODUCTION

Alzheimer’s disease (AD) is a debilitating neurodegenerative disease, characterized by an array of cognitive defects including impairments in memory, executive function, language, and sensation (1). Over time, the disease progresses to severe dementia and eventually leads to death. Approximately 6.9 million Americans have AD and there is no treatment to prevent or reverse the disease (2).

One of the hallmarks of AD is hyperphosphorylation of tau proteins which causes the formation of intracellular neurofibrillary tangles (NFTs) (3). Tau is a microtubule-associated protein that aids in assembling and stabilizing microtubules. This, in turn, facilitates axonal transport and maintains neuronal function and network (4).

Hyperphosphorylation of tau interrupts its binding to microtubules, causing misfolding and aggregation in the form of NFTs, which results in microtubule instability and neuroinflammation, which eventually leads to neurodegeneration in not only AD but other forms of tauopathies as well (5).

While AD has been viewed as a brain disease, recent studies have found the expression of AD-related protein aggregates in other organs, including the heart (6, 7). Emerging evidence indicates tau is expressed in normal cardiomyocytes (8, 9); and mice deficient in tau developed age-dependent cardiovascular dysfunction and cardiac hypertrophy (9), suggesting an important role for tau in cardiac physiology.

Phosphorylated tau accumulation and tau aggregates have also been identified in both brain and heart samples of AD humans (8), which highlights the systemic nature of tau pathology and demands a better understanding of the potentially shared pathogenic mechanisms between the neural and cardiac pathologies of AD.

Growing evidence suggests that insufficient clearance of misfolded proteins (e.g., tau and amyloid beta) plays an important role in most cases of AD (10). Therefore, investigating the roles of protein degradation pathways in AD may reveal new insights into AD pathogenic mechanisms and potential treatment approaches. The ubiquitin-proteasome system (UPS) is the primary mechanism for degrading terminally misfolded proteins in the cell, thereby pivotal to protein quality control (11). Most, if not all, misfolded proteins are degraded by sequential steps through the UPS (12). Multiple studies have revealed a connection between AD tauopathy and UPS dysfunction (13, 14). Tau is physically associated with the proteasome in human AD brain cells and aberrant tau reduces proteasome activities (15, 16). Activating the proteasome by cAMP augmentation was associated with a reduction of tau accumulation and an amelioration of cognitive impairment in a tauopathy mouse model (16). Phosphorylation of the proteasome is a critical mechanism for regulating 26S proteasome activities (17, 18), which can be achieved by activating protein kinases such as the cAMP-dependent protein kinase or protein kinase A (PKA) and protein kinase G (19). Both *in vitro* and *in vivo* studies have established that phosphorylation of the 19S subunit, RPN6 (regulatory particle non-ATPase 6) at Serine 14 (p-S14-RPN6) is induced specifically by PKA and mediates PKA-induced proteasome activation (20–22). However, the pathophysiological significance of p-S14-RPN6 in AD remains to be established.

To address these critical gaps, the current study was conducted to determine the role of homeostatic p-S14-RPN6 or, by extension, the role of PKA-mediated activation of 26S proteasomes in the progression of brain and cardiac pathology of a tauopathy-based AD mouse model, where our recently created and validated Rpn6^S14A^ knock-in mice were employed to block genetically p-S14-Rpn6 (22). Our findings provide compelling evidence that homeostatic p-S14-RPN6 or the activation of 26S proteasomes by PKA protects against the perturbation of proteostasis and attenuates pathologies in both the brain and heart of the tauopathy mice. The findings also imply that the augmentation of p-S14-RPN6 may be a key mechanism underlying the therapeutic effects of stimulating cAMP/PKA in treating tauopathy-based AD, and potentially other neurodegenerative disorders.

## RESULTS

### Genetic blockade of p-S14-Rpn6 exacerbated the decline of cognitive and motor functions in PS19 mice

We recently created and validated the Rpn6^S14A^ mouse, in which the endogenous *Rpn6/Psmd11* gene is modified such that the codon for Serine 14 of Rpn6 is mutated to code for Alanine to block phosphorylation (22). The homozygous Rpn6^S14A^ (referred to as S14A hereafter) mice are viable and fertile and do not display gross abnormality or cardiac malfunction within 1-year of age (23). The PS19 mouse (RRID: IMSR_JAX:008169) has been widely used as a tauopathy and AD model (24). To investigate the role of homeostatic p-S14-RPN6 in AD progression, we introduced the S14A allele into the PS19S mice via multiple rounds of crossbreeding; the resultant littermate wild type (WT), PS19, and PS19 coupled with homozygous S14A (PS19::S14A/A) mice were used for studies reported here. To determine the protein levels of Ser14-phosphorylated Rpn6 (pS14-Rpn6) in PS19 mice and confirm the genetic blockade of p-S14-Rpn6 in PS19::S14A/A mice, we first performed western blot analyses for myocardial pS14-Rpn6 and total Rpn6 in littermates at 3 months of age (3m). As expected, pS14-Rpn6 was completely absent in PS19::S14A/A mice, whereas its expression was not significantly different between WT and PS19 mice (**Figure 1A, 1B**). A statistically significant increase in the total Rpn6 level was detected in both PS19 and PS19::S14A/A mice (**Figure 1A, 1C**), suggesting that the proportion of pS14-Rpn6 is reduced in PS19 mice compared to WT mice.

**Figure 1.**
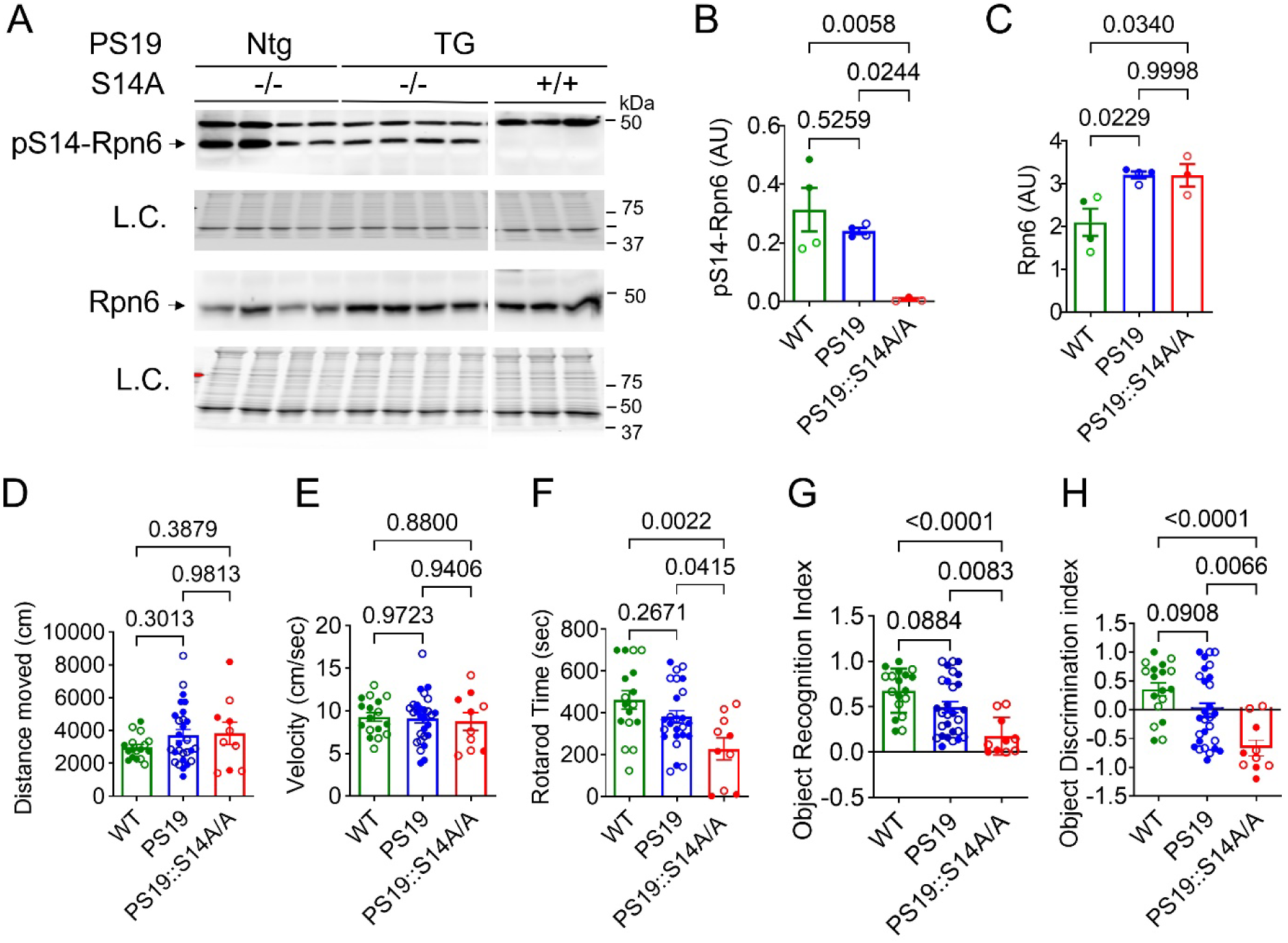
Effect of Rpn6^S14A^ on cognitive and motor function of PS19 mice. (**A**-**C**) Crude protein extracts from the ventricular myocardium of wild type (WT) (i.e., PS19 Ntg::S14A-/-), PS19 (PS19 TG::S14A-/-), and PS19::S14A/A (PS19 TG::S14A+/+) mice at 3 months, were subjected to an SDS-PAGE followed by immunoblot analyses for the indicated proteins. Representative images (**A**) and pooled densitometry data (**B**-**C**) of western blot analyses for Ser14-phosphorylated Rpn6 (pS14-Rpn6) and total Rpn6. Each lane or dot represents an individual mouse. L.C. indicates loading control, showing a segment of the in-lane total protein signal from the stain-free total protein imaging employed to normalize the loading; the same method is also used in all other western blots. (**D**-**H**) Neurobehavioral tests were performed on WT (n=17; 9 males, 8 females), PS19 (n=28; 15 males, 13 females), and PS19::S14A/A (n=10; 4 males, 6 females) mice at 6 months of age. (**D** and **E**) Total distance travelled (**D**) and mouse velocity (**E**) from the open field test. (**F**) Time spent on Rotarod. (**G** and **H**) Object recognition index (**G**) and object discrimination index (**H**) from the novel object test. Scatter dot plot superimposed by mean±SEM; each dot corresponds to an individual mouse; solid and open symbols denote males and females, respectively; the same convention applies to other figures as well. Shown above brackets are *P* values derived from One-way ANOVA followed by Tukey’s test.

In PS19 mice, cognitive decline is one of the most prominent features of AD (25). Therefore, we performed behavioral tests to characterize the changes in cognitive processes resulting from p-S14-Rpn6 blockage in 6m-old PS19 mice. We employed an open field test (OFT) to analyze exploratory behavior and general activity. We observed that travel distances and velocity were comparable among the groups (**Figures 1D and 1E**), which suggests that genetic blockade of p-S14-Rpn6 does not affect the general locomotor activity of the PS19 mice. Previous research has indicated that PS19 mice also exhibit progressive motor impairment and muscle atrophy starting at 6m of age (26). We therefore used the rotarod test to determine the impact of loss of p-S14-Rpn6 on motor function and coordination abilities. The PS19::S14A/A group showed a significantly shorter time duration to stay on the rotating rotarod than PS19 (*P*=0.04) and WT (*P*=0.0022) although the difference in the rotarod time between PS19 and WT mice did not reach statistical significance at this age (**Figure 1F**), indicating that loss of p-S14-Rpn6 expedites the impairment of motor function and coordination abilities in mice with tauopathy. To observe short-term memory impairment, we used a novel object recognition test. PS19::S14A/A mice displayed exacerbated cognitive dysfunction as indicated by marked reduction of both the object recognition index and the object discrimination index, compared to WT (*P*<0.0001) and PS19 (*P*=0.0083; 0.0066) mice (**Figures 1G and 1H**). These findings thus provide compelling evidence that genetic blockade of p-S14-Rpn6 or, by extension, loss of the PKA-mediated activation of 26S proteasomes aggravates cognitive and motor deficits in the AD mice.

### Genetic blockade of p-S-14-Rpn6 expedited and exacerbated cardiac malfunction of PS19 mice

Recent research has revealed that AD tauopathy is a multifactorial disease and affects not only the brain but the heart as well (6, 7). Therefore, to test the effect of blocking p-S14-Rpn6 on cardiac function, we performed 2D-guided M mode echocardiography on WT, PS19, and PS19::S14A/A mice at 3m, 6m, and 9m (see **Figure S1A** for representative images). Normally, heart size and cardiac output (CO) are proportional to body size; hence, we normalized the cardiac morphometric parameters to body weight (BW), to improve the comparability among different genotypes. All three genotype groups had comparable mouse BWs at 3m and 6m, but PS19::S14A/A mice were significantly lighter than WT at 9m (**Figure S2A**, *P*=0.048), consistent with skeletal muscle atrophy. At all 3 times points, no statistically significant differences in BW-adjusted LV end-diastolic internal diameter (LVID;d) and LV end-diastolic volume (LVEDV;d) were detected among all 3 genotype groups (**Figure 2A**, **2B**), but the fractional shortening (FS) and ejection fraction (EF) of both PS19 and PS19::S14A/A groups began to decline at 3m compared with the WT group (**Figure 2C**, **2D**), with the declines in the PS19::S14A/A group but not the PS19 group reaching statistical significance (*P*=0.012, 0.017, respectively). By 6m, the declines of FS and EF in both PS19 and PS19::S14A/A group became statistically significant and, by 9m, the PS19::S14A/A group showed more pronounced declines than the PS19 group (*P*=0.0033, 0.0031, respectively). A similar pattern of temporal changes in BW-adjusted stroke volume (SV) and cardiac index (CO/BW) was also observed among the three groups (**Figure 2E**, **2F**). These data show that the impairment of cardiac systolic function in the AD mice occurs earlier and deteriorates faster when p-S14-RPN6 is blocked.

**Figure 2.**
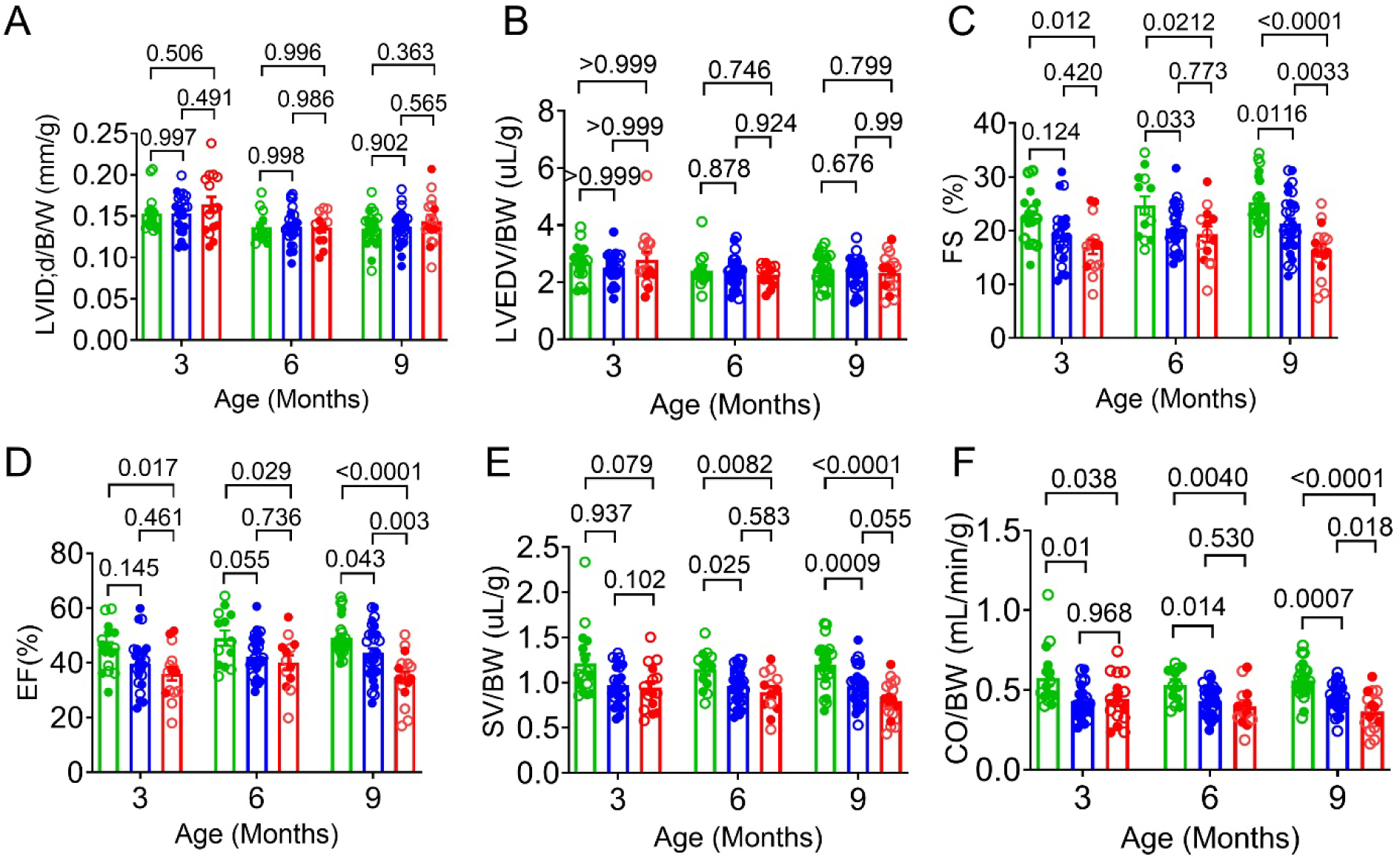
The effect of Rpn6^S14A^ on tau-induced cardiac malfunction. Mice of the indicated genotypes (WT, green; PS19, blue; PS19::S14A/A, red) were subjected to echocardiography at the indicated age. (**A**) LV end-diastolic internal diameter (LVID;d) to body weight (BW) ratio. (**B**) LV end-diastolic volume (LVEDV) to BW ratio. (**C**) Fractional shortening (FS). (**D**) Ejection fraction (EF). ((**E**) Stroke volume (SV) to BW ratio. (**F**) Cardiac output (CO) to BW ratio. For WT mice, n=9 males+7 females at 3m, n=7 males+6 females at 6m, and n=11 males+14 females at 9m; for PS19 mice, n=14 males+9 females at 3m, n=15 males+15 females at 6m, and n=15 males+14 females at 9m; for PS19::S14A/A mice, n=5 males+10 females at 3m, n=7 males+6 females at 6m, and n=6 males+11 females at 9m. Scatter dot plot superimposed by mean±SEM; each dot corresponds to an individual mouse. Shown above brackets are *P* values derived from One-way ANOVA followed by Tukey’s test, except for LVEDV/BW at 3m where Kruskal Wallis followed by Dunn’s test was applied because of non-normally distributed data.

To monitor LV diastolic function, we recorded and analyzed tissue Doppler at the level of the mitral valve annulus as well as pulsed-wave Doppler of the mitral valve at 3m and 6m (**Figures S3** and **S4**). Although no statistically significant differences were detected at 3m, we observed a significant increase in the ratio of early diastolic inflow and septal annulus velocity (E/e’) in PS19::S14A/A mice as compared to WT (*P*=0.0004) and PS19 (*P*=0.014) at 6m (**Figure S4A**). Increased E/e’ ratio is a strong indicator of diastolic dysfunction which typically indicates impaired left ventricular relaxation (27). Thus, these data suggest that blockade of p-S14-RPN6 also impairs diastolic function of the heart of AD mice, but this impairment appears to occur later than systolic impairment. Taken together, these findings suggest that the genetic blockade of p-S14-Rpn6 leads to progressive deterioration of both systolic and diastolic function of the heart in PS19 mice.

### Genetic blockade of p-14-Rpn6 exacerbated hippocampal gliosis and microglial activation in PS19 mice

In addition to the cognitive and motor function deficits, the PS19 mouse model exhibits gliosis and microglial activation as a neuroinflammatory response to tau accumulation at 3m of age (24). Hence, we investigated the potential effects of p-S14-Rpn6 blockade on tau-induced gliosis and microglial activation in the brain using GFAP (glial fibrillary acidic protein) as an astrocyte marker and IBA1 (an ionized calcium-binding adaptor molecule 1) as a microglial marker. Our western blot analyses detected increased hippocampal expression of GFAP and IBA1 proteins in PS19::S14A/A mice as compared to WT (*P*<0.0001; *P*=0.0029) and PS19 mice (*P*=0.0018; *P*=0.049) (**Figures 3A, 3B, 3E, 3F**). To further validate our results, immunofluorescence microscopy was performed to examine astrocytes and microglia in the coronal sections of the brain. Both GFAP (**Figure 3C, 3D**) and IBA1 (**Figure 3G, 3H**) showed increased immunoreactivity in the PS19::S14A/A mouse hippocampi than in the WT (*P*<0.0001; *P*<0.0001) and the PS19 (*P<0.0001*; *P*=0.0008) mice, which is indicative of increased astrogliosis and microglial activation in the PS19::S14A/A mice. Collectively, these data reveal that blockage of p-S14-Rpn6 augments inflammatory responses and gliosis in the brain of the AD mice.

**Figure 3.**
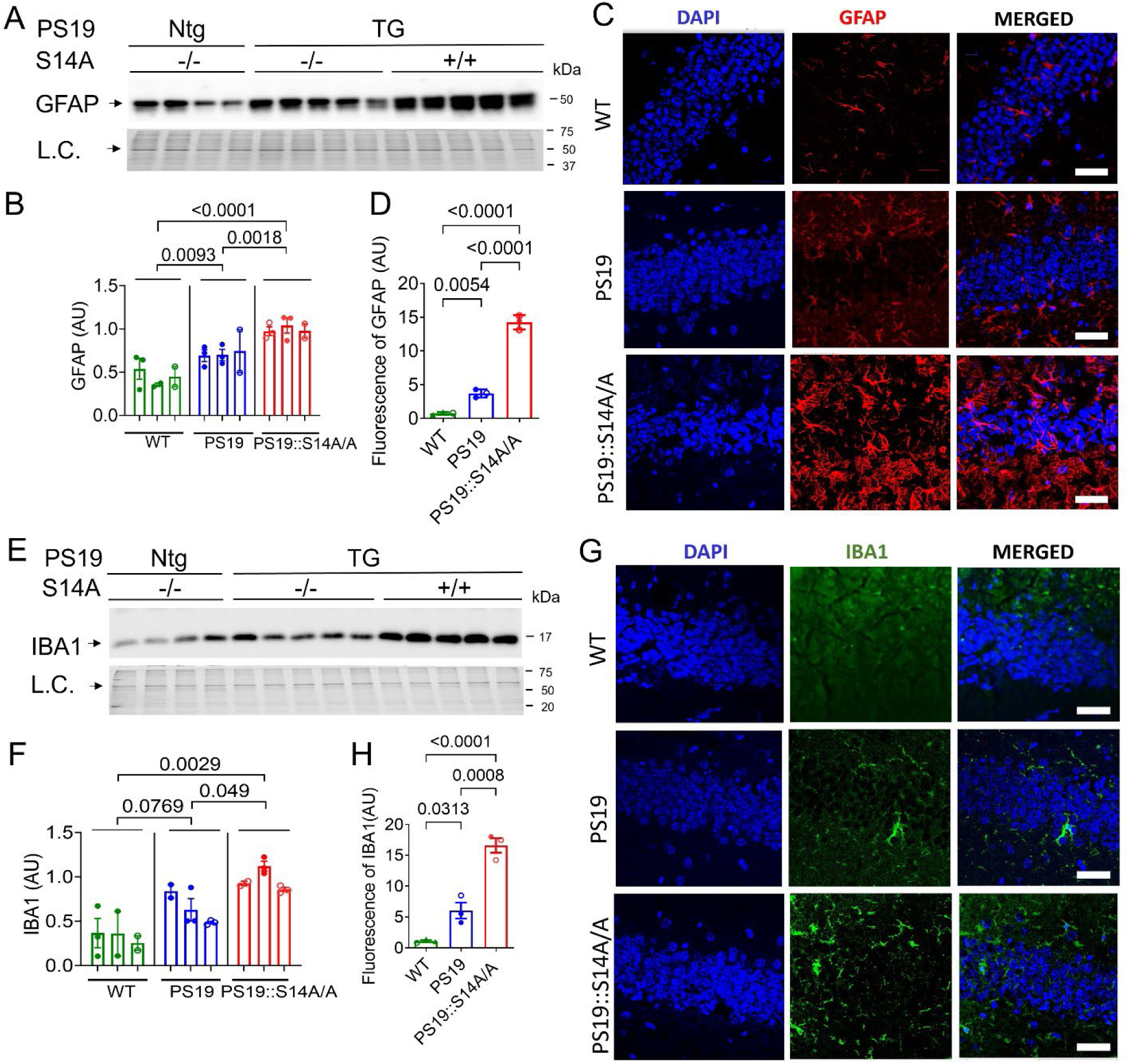
Effect of Rpn6^S14A^ on the inflammatory response in the brain of PS19 mice. Crude protein extracts from the hippocampus of wild type (WT, Ntg::S14A-/-), PS19 (PS19 TG::S14A-/-), and PS19::S14A/A (PS19 TG::S14A+/+) mice at 9 months of age were subjected to SDS-PAGE followed by immunoblot analyses for the indicated proteins (**A, B, E,** and **F**), and fixed brain tissue sections for immunohistochemistry (**C**, **D**, **G**, and **H**). (**A** and **E**) Representative images and (**B** and **F**) pooled densitometry data of western blot analyses for GFAP (glial fibrillary acidic protein) and IBA1(Ionized calcium-binding adaptor molecule 1), respectively. Scatter dot plot superimposed by mean±SEM; each bar corresponds to an individual mouse (n=3 mice for each group) and each dot represents a technical repeat. Shown above brackets (B, F) are *P* values derived from nested one-way ANOVA followed by Tukey’s test with technical replicates nested within biological replicates. (**C** and **G**) Representative confocal micrographs and (**D** and **H**) quantitative analysis of immunofluorescence staining for GFAP and IBA1, respectively. Nuclei are stained blue with DAPI. Scale bar=25 μm; n=3 mice per group. Shown above the brackets (D, H) are *P* values derived from one-way ANOVA followed by Turney’s tests.

### Genetic blockade of p-14-Rpn6 augmented synaptic and neuronal losses in the hippocampus of PS19 mice

Several studies have suggested that hyperphosphorylation, mis-localization, or overexpression of tau induce toxicity by impairing synaptic function in AD (28, 29). To investigate the effect of blockade of p-S14-Rpn6 on synaptic integrity, we performed western blot analyses for synaptic-associated proteins (PSD-95 and synaptotagmin) in the hippocampus at 9m. We detected a substantial reduction in synaptotagmin (*P*=0.0005; P=0.029) and PSD-95 (*P*=0.008; *P*=0.068) in PS19::S14A/A mice, compared with WT and PS19 littermates (**Figures 4A ∼ 4C**). To further locate these changes, we performed double immunofluorescence labeling for PSD-95 and synaptotagmin in the dentate gyrus of the hippocampus. Consistent with our western blot results, PS19::S14A/A mice displayed significantly reduced immunoreactivity for PSD-95 (green) and synaptotagmin (red) in the hippocampal region when compared to the WT (*P*=0.0020; *P*<0.0001) and the PS19 (*P*=0.048, P=0.028) groups (**Figure 4E, 4F**).

**Figure 4.**
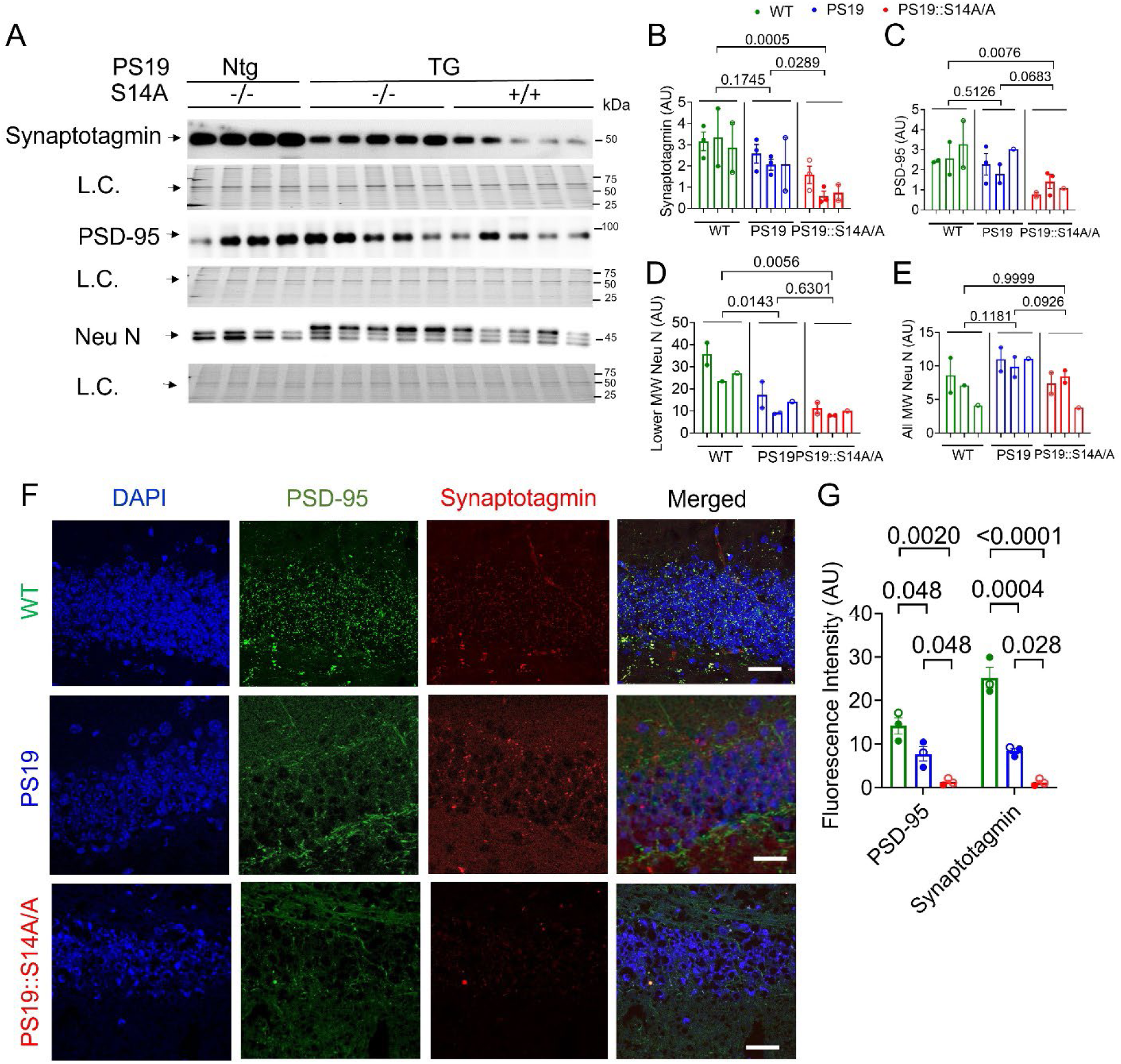
Effect of Rpn6^S14A^ on hippocampal synapsis and neuron loss in PS19 mice. (**A**-**D**) Crude protein extracts from the hippocampus of wild type (WT, Ntg::S14A-/-), PS19 (PS19 TG::S14A-/-), and PS19::S14A/A (PS19 TG:: S14A+/+) mice at 9 months were subjected to SDS-PAGE followed by immunoblot analyses for the indicated proteins. Representative images (**A**) and pooled densitometry data (**B**-**E**) of western blot analyses for synaptotagmin (**B**), postsynaptic density protein-95 (PSD-95) (**C**), and neuronal nuclear protein (Neu N) (**D, E**). For quantitative data of Neu N, panel D shows densitometric data of the two lower <u>m</u>olecular <u>w</u>eight (MW) bands seen in all 3 groups, wheras panel E presents data from all three MW bands. Scatter dot plot superimposed by mean±SEM; each bar represents an individual mouse (n=3 mice/group) and each dot corresponds to a technical repeat. Shown above brackets (B-E) are *P* values derived from Nested One-way ANOVA followed by Tukey’s test with technical replicates nested within biological replicates. (**F**) Representative confocal micrographs and (**G**) quantitative analysis of double immunofluorescence staining of PSD-95 and synaptotagmin in the 5 μm coronal sections of the indicated genotype. Scale bar=25 μm; n=3 mice/group. *P* values shown above brackets (F) are derived from One-way ANOVA followed by Tukey’s test.

Reduction in these synaptic proteins further prompted us to look for an overall loss of neurons in PS19 mice with blockade of p-S14-Rpn6. Thus, we performed western blot analyses for Neu N (**Figure 4A, 4D**), a neuronal nuclear protein that is often used as a marker of postmitotic neurons (30). We detected that the higher molecular weight band (∼48kDA) was clearly increased, but the level of the two lower molecular weight bands (∼45kDa) were significantly reduced in the PS19 and the PS19::S14A/A mice as compared to WT (**Figure 4D**; *P*=0.014, 0.006); however, no significant difference between PS19 and PS19::S14A/A mice (*P*=0.630). When all three bands of Neu N were quantified together, the level of Neu N in PS19::S14A/A tended to be lower than in the PS19 group (*p*=0.093; **Figure 4E**). Overall, the above data provide compelling evidence that blocking p-S14-Rpn6 exaggerates tauopathy associated hippocampal synaptic and neuronal losses in PS19 mice.

### Genetic blockage of p-S14-Rpn6 expedited cardiac pathology in PS19 mice

The echocardiographic data presented in Figure 2 indicate that blocking p-S14-Rpn6 is progressively detrimental to cardiac function. Echocardiography also revealed a significantly increased LV posterior wall thickness at the end diastole normalized to BW (LVPW;d/BW) in both PS19 and PS19::S1A/A (**Figure S1H**, P=0.014, P=0.001), suggesting that cardiac hypertrophy and remolding occur in PS19 mouse hearts. This prompted us to further examine cardiac pathology at the gravimetric, histological and molecular levels. Our gravimetric assessment confirmed the finding of cardiac hypertrophy with significantly increased heart weight (HW) to BW (**Figure 5D**; *P*=0.012) and ventricular weight (VW) to BW ratio (**Figure 5E**; *P*=0.0193) in PS19::S14A/A mice but not PS19 mice (*P*=0.061, 0.180), as compared to the WT group. In addition, PS19::S14A/A mice but not PS19 mice had significantly increased lung weight to body weight ratio (**Figure 5G**; *P*=0.040) as compared to WT, which is another indicator of left-sided heart failure. Therefore, to get further insight into cardiac pathology, we decided to investigate cardiac remodeling in the PS19 mouse.

**Figure 5.**
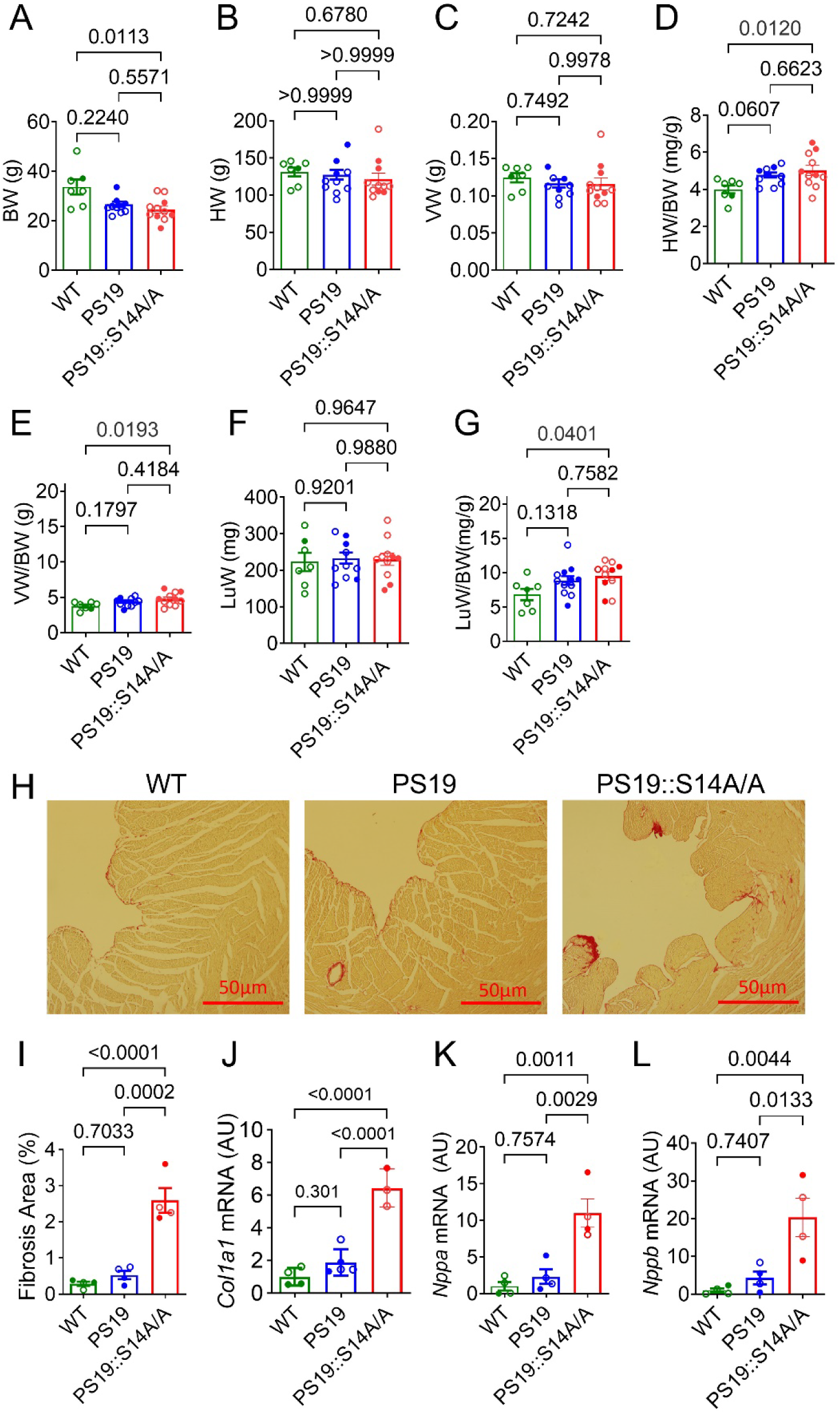
Effect of Rpn6^S14A^ on cardiac remodeling and fetal gene program of PS19 mice. (**A**-**G**) Gravimetry data of WT, PS19, and PS19::S14A/A mice at 9 months. (**A**) Body weight (BW). (**B**) Heart weight (HW). (**C**) Ventricle weight (VW). (**D**) HW to BW ratio. (**E**) VW

During pathological cardiac remodeling, excessive extracellular matrix (ECM) deposition occurs, leading to myocardial fibrosis. Excessive accumulation of ECM proteins, especially collagen I and III are the hallmarks of cardiac fibrosis (31). Therefore, we performed picrosirius red staining, which revealed increased collagen staining in PS19::S14A/A relative to WT (*P*=<0.0001) and PS19 (*P*=<0.0001) mice (**Figure 5H** and **5I**). These results were further validated by ventricular mRNA expression of collagen I (Col1a1), where we observed significantly upregulated mRNA expression of *Col1a1* (**Figure 5J**) in PS19::S14A/A mice as compared to PS19 (*P*=0.0058) and WT (*P*=0.0004) mice. During cardiac remodeling, cardiac stress leads to reactivation of the fetal gene program; for example, upregulation of genes encoding atrial and brain natriuretic peptides (*NPPA* and *NPPB*, respectively) in the ventricular myocardium is often associated with cardiac pathology (32), which even is a prognostic indicator of clinical severity (33). Therefore, we determined the mRNA levels of *Nppa* and *Nppb* in ventricular myocardium (**Figure 5K, 5L**).

As anticipated, our PS19::S14A/A mice displayed significantly higher mRNA expression of both *Nppa* and *Nppb* as compared to WT (*P*=0.001; *P*=0.004) and PS19 mice (*P*=0.003; *P*=0.013). These results confirm that at 9m of age, these PS19::S14A/A mice exhibit pathophysiological changes indicative of a progression toward heart failure. Overall, the data presented here indicate that the genetic blockade of p-S14-Rpn6 exacerbates cardiac pathology in a tauopathy-based AD.

### Genetic blockade of p-14-Rpn6 exacerbated tau pathology in the brain and heart of PS19 mice

Intraneuronal accumulation of aberrant forms of tau is a hallmark of AD (34). Hence, to determine whether suppressing proteasome activity affects tau pathology in the brain and heart of PS19 mice, we first measured total tau (both isoforms of mouse and human origin) in hippocampal and myocardial tissue using the A10 antibody. At 9m, S14A/A coupled PS19 mice displayed significant increases in total tau proteins in both hippocampal and myocardial tissues (**Figure 6A, 6B, 6F, 6G**), as compared to PS19 (*P*=0.0049; *P*=0.0027) and WT (*P*=0.0008; *P*=0.0001) mice.

**Figure 6:**
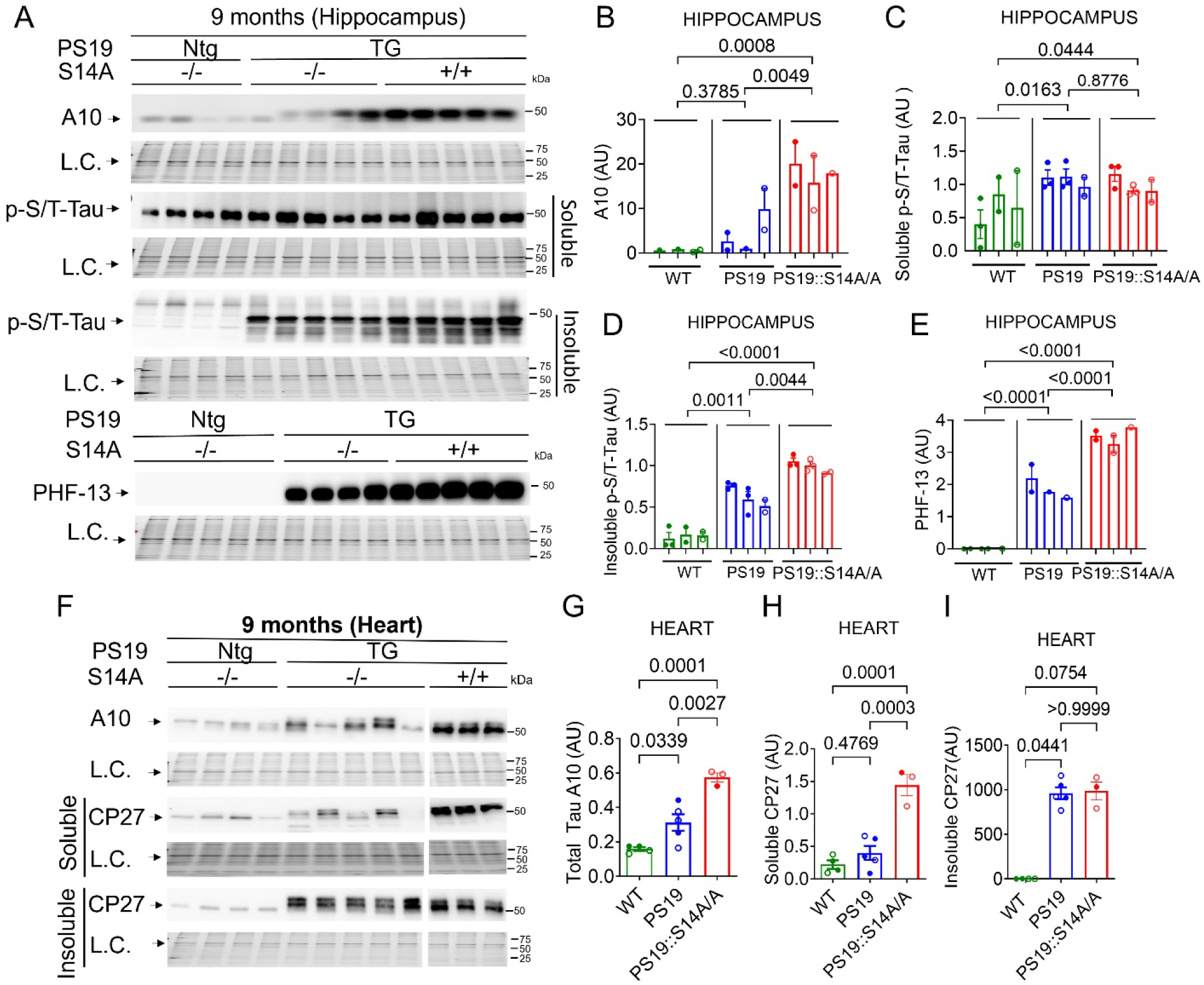
Rpn6^S14A^ exacerbated tau pathology in the brain and heart of PS19 mice. Crude protein extracts from the brain hippocampus (**A**-**E**) and ventricular myocardium (**F**-**I**) of wild type (WT; PS19 Ntg::S14A-/-), PS19 (PS19 TG::S14A-/-), and PS19::S14A/A (PS19 TG:: S14A+/+) mice at 9 months were subjected to SDS-PAGE followed by immunoblot analyses for the indicated proteins. (**A**-**E**) Representative images (**A**) and pooled densitometry data (**B**-**E**) of western blot analyses for total tau proteins that are reactive to the A10 antibodies (A10) (**B**), Ser202/Thr205-phopshorylated tau (p-S/T-Tau) in the NP40 soluble and insoluble fractions (**C** and **D**), and PHF-13 antibodies detected total tau proteins (**E**) in mouse hippocampus. Each bar represents an individual mouse, and each dot corresponds to a technical repeat. Shown above brackets are *P* values derived from Nested One-way ANOVA followed by Tukey’s test with technical replicates nested within biological replicates. (**F**-**I**) Representative images (**F**) and pooled densitometry data (**G**-**I**) of western blot analyses for myocardial tau proteins using the A10 antibodies (A10) (**G**) or the CP27 antibodies (NP40 soluble and insoluble fractions) (**H** and **I**). Scatter dot plot superimposed by mean±SEM; each dot corresponds to an individual mouse (n = 3-5 mice per group). Shown above brackets are *P* values derived from One-way ANOVA followed by Tukey’s test, except for CP27 insoluble fraction where Kruskal Wallis followed by Dunn’s test was used because of non-normally distributed data.

Among many forms of tau, hyperphosphorylated tau is the major component of the paired helical filament (PHF) of AD (35). Therefore, to get further insight into tau pathology, we probed for Ser202/Thr205-phosphorylated tau (p-S202/T205-Tau) in the NP40 (Nonidet P40) substitute (nonylphenylpolyethylene glycol) soluble and insoluble fractions of hippocampal tissue lysates and for PHF-13 in total hippocampal tissue lysates. As illustrated in **Figure 6A** and **6C**, p-S202/T205-Tau was elevated in both PS19 and PS19::S14A/A mice as compared to the WT brain (*P*=0.0163, 0.04) but did not show a significant difference between the two PS19 groups (*P*=0.8776) in the soluble fraction of the hippocampus. However, we observed that the increases of p-S202/T205-Tau in the insoluble fraction were significantly greater in PS19::S14A/A compared to PS19 (P=0.0044) (**Figure 6D**), which indicates that aberrant protein aggregation increases when p-S14-Rpn6 is blocked in PS19 mice. Further, as detected with the PHF13 antibody, western blot analysis revealed a significantly higher accumulation of Ser396-phosphorylated tau in PS19::S14A/A mice as compared to PS19 (*P*=<0.0001) and showed no expression in WT hippocampi (*P*<0.0001) (**Figure 6E**).

As previously described, the PS19 mouse line overexpresses the human mutant form of tau (P301S), with a 3-5-fold greater expression of human tau compared to the naturally occurring mouse tau (24). Therefore, in the next set of experiments, we attempted to determine the myocardial expression of human tau using the CP27 antibody (**Figure 6F-6I**). We found that PS19::S14A/A animals exhibited a significantly higher level of the CP27-reactive tau in in the soluble fraction of myocardial tissue than PS19 (*P*=0.0003) and WT (*P*=0.0001) mice; however, in the insoluble fraction, the increases of C27-reactive tau in the PS19 and in the PS19::S14A/A mice over WT mice are comparable (*P*>0.9999).

Taken together, these findings reveal that in PS19 mice, there is an increased accumulation of total and humanized tau, and genetic blockade of p-S14-Rpn6 exacerbates the accumulation of these pathological tau and the progression of tau pathology in not only the brain but also the heart. Additionally, these findings highlight the interconnected nature of tau pathology in different organs and emphasize the need for further research into the cardiac pathology and mechanisms underlying this phenomenon.

### Genetic blockade of p-S14-Rpn6 impaired the proteostasis in PS19 mice

Research has shown that aberrant protein aggregation and Ub conjugate levels increase when there is a diminished proteasome activity (36). Therefore, we assessed changes in hippocampal and myocardial Ub conjugate levels in PS19::S14A/A, PS19, and WT mice (**Figure 7**). PS19::S14A/A mice displayed considerably higher expression of hippocampal and myocardial Ub conjugates than PS19 (*P*=0.046, *P*=0.0064;) and WT mice (P=0.0164, P=0.0006), indicating that the transgenic tau induced impairment of proteostasis in both the brain and heart was exacerbated by loss of p-S14-Rpn6. Since ubiquitinated proteins are the substrates of the 26S proteasome, elevated Ub conjugates in the brain and heart of PS19::S14A/A mice are consistent with the notion that homeostatic p-S14-Rpn6 play a key compensatory role in promoting proteasome-mediated protein degradation in the AD mice.

**Figure 7.**
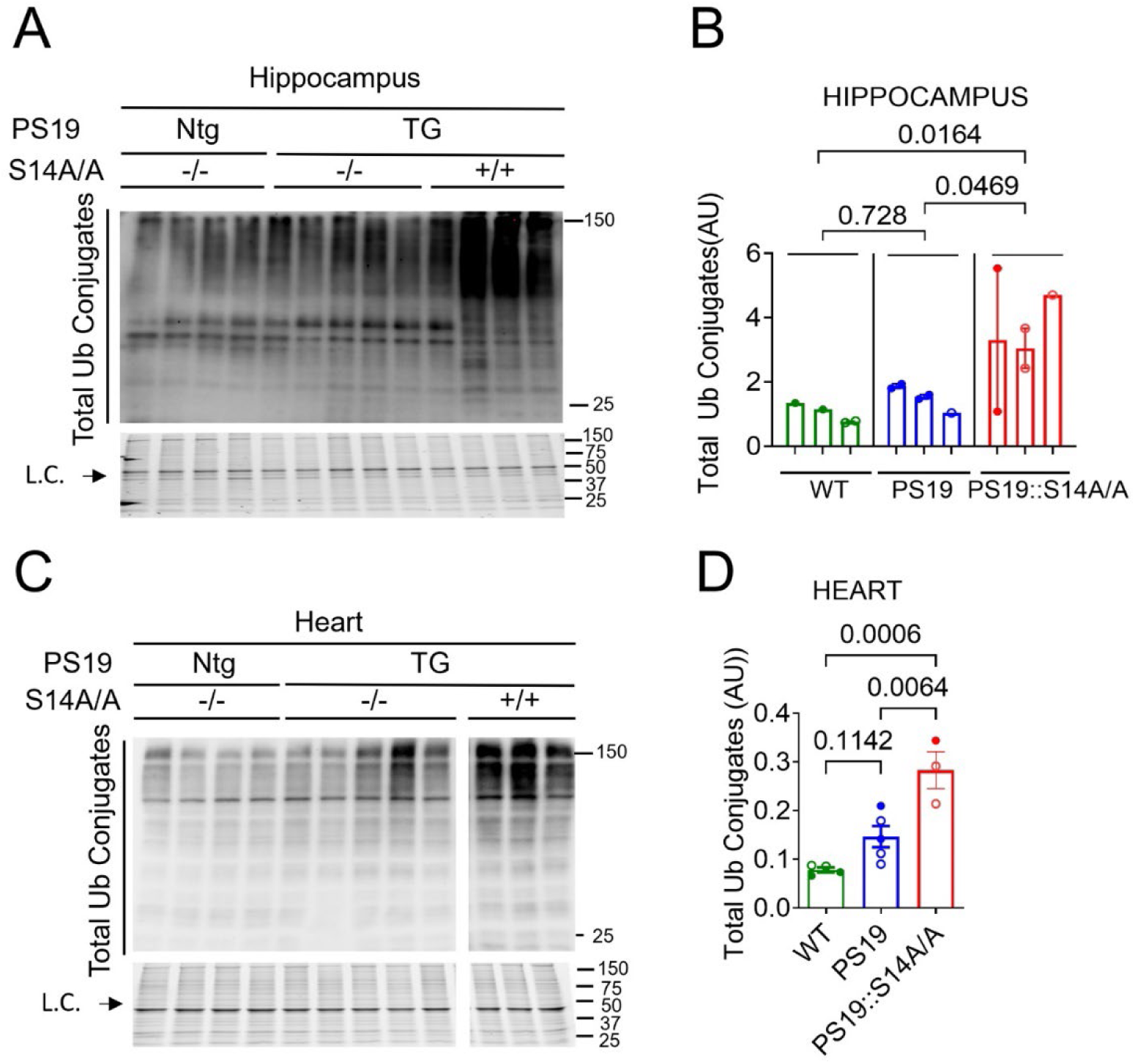
Effect of Rpn6^S14A^ on proteostasis of PS19 mouse brains and hearts. Crude protein extracts from the brain hippocampus (**A**, **B**) and ventricular myocardium (**C**, **D**) of wild type (WT; PS19 Ntg::S14A-/-), PS19 (PS19 TG::S14A-/-), and PS19::S14A/A (PS19 TG:: S14A+/+) mice at 9 months of age were subjected to SDS-PAGE followed by immunoblot analyses for the indicated proteins. (**A** and **B**) Representative images (**A**) and pooled densitometry data (**B**) of western blot analyses for total ubiquitin (Ub) conjugates in the hippocampus. Each bar represents an individual mouse, and each dot corresponds to a technical repeat. Shown above brackets are *P* values derived from nested one-way ANOVA followed by Tukey’s test with technical replicates nested within biological replicates. (**C** and **D**) Representative images (**C**) and pooled densitometry data (**D**) of western blot analyses for total Ub conjugates in the heart. Each lane or dot corresponds to an individual mouse (n = 3-5 mice per group). Shown above brackets are *P* values derived from one-way ANOVA followed by Tukey’s test. Scatter dot plot superimposed by mean±SEM.

## DISCUSSION

In the present study, we have demonstrated that the genetic blockade of p-S14-Rpn6 aggravates cognitive and motor function deficits and deteriorates both the systolic and diastolic function of the heart in PS19 tauopathy mice. We have further shown that the blockade of p-S14-Rpn6 exaggerates neuroinflammatory responses and reduces synaptic integrity in the PS19 mouse brain. In addition, various findings related to tau pathology indicate that the blockade of p-S14-Rpn6 also exacerbates tau pathology in the myocardium of PS19 mice. More pronounced disrupted proteostasis was also evident in PS19::S14A/A mouse hearts and hippocampi as shown by the increases in hippocampal and myocardial ubiquitinated proteins in PS19::S14A/A mice. These new findings provide compelling evidence that homeostatic p-S14-Rpn6 and, by extension, the activation of 26S proteasomes by PKA plays a major protective role against tauopathy-based AD disease progression; accordingly, it is implicated that increasing p-S14-Rpn6 through for example augmentation of cAMP/PKA may represent potentially a new therapeutic strategy to slow down the progression of both heart and brain-related pathologies in tauopathy based AD.

The PS19 mouse model used in our study overexpresses the human mutated form of tau (MAPT (1N4R) tau P301S) driven by the mouse prion promoter and shows 5 times higher human tau expression than that of endogenous mouse tau (24).

Hyperphosphorylated tau in these mice forms neurofibrillary tangles (NFTs), serves as damage-associated molecular patterns (DAMPs), and activates toll-like receptors (TLRs) of glial cells (37), which in turn change the morphology of normal astrocytes to neurotoxic form, termed gliosis. The upregulated gliosis and microglial activation, in turn, causes the release of proinflammatory cytokines, such as TNFα, and IL-1β, which contributes to synaptic and neuronal loss mediated by activating pro-apoptotic factors and caspase-3 (38, 39), and ultimately leads to brain atrophy and cognitive decline.

Research related to tauopathy pathology in the heart of PS19 is relatively scarce. None of the current research has directly identified the mechanism of cardiac dysfunction in AD especially in tauopathy mice (PS19). However, previous studies have linked heart failure and AD in humans (40). Another study has demonstrated that there is an elevated Aβ in the circulation and thus in cardiomyocytes, which disrupts mitochondrial respiration and in turn, increases oxidative stress and inflammation, which leads to reduced systolic function, and cardiac fibrosis in the 5XFAD model (41). Luciani, *et al*. have demonstrated that highly toxic tau oligomers are present in human tau mice and human myocardium, which alters microtubule tyrosination and detyrosination balance, required for cargo trafficking, including Ca2+ handling proteins, leads to increased myocardial stiffness, diastolic dysfunction, and consequently heart failure. However, the study did not observe a significant decline in systolic function until 12 months of age (8). In contrast to the existing studies, we observed for the first time to our best knowledge, that a tau-induced decline in cardiac index starts from 3 months of age in the PS19 mice, which further declines in an age-dependent manner; however, the Doppler-derived parameters of diastolic malfunction did not reach statistical significance in the PS19 mice during the first 6m (Figures 2 and S4). This discrepancy could be because of the difference in the mouse line, or a smaller number of mice used by the prior study (8). Thus, we speculate that tau may play a role in cardiac pathogenesis in PS19 mice, either because the mouse prion promoter, which drives overexpression of the human mutant form of tau in PS19 mice, is actively expressed in the heart as well (42), which itself might be causing increased expression of tau in the heart and contributing to tau-related pathological changes; or alternatively it is because of an inter-organ migration of the pathogenic tau between the brain and heart, caused by disrupted blood-brain barrier in AD (43), leading to increase circulatory tau in blood and thereby in the heart. Evangelisti and colleagues (7) have suggested the same that either tau migrates from organ to organ metastatically, or it is a systemic disease that has a similar mechanism arising in multiple organs independently. Relatedly, there is a growing list of organs affected by tauopathy (44, 45). However, the mechanism of tauopathies in other organs, including the heart of PS19 remains unexplored.

UPS dysfunction has been implicated, as a primary cause or a secondary consequence, in the neural pathogenesis of AD. The 26S proteasome is primarily responsible for degrading misfolded or damaged proteins (46). These misfolded proteins i.e. hyperphosphorylated tau, bind with the 26S particle but resist degradation or the ATPases translocation, thereby disrupting the multistep process of degradation of other proteins, leading to the accumulation of toxic aggregates in neurons and systemic organs, including the heart (43).

The current study has demonstrated that p-S14-Rpn6 plays a critical role in response to misfolded tau in the heart and brain. In our experiments, the PS19::S14A/A mice displayed more elevated accumulation of tau in both hippocampus and heart compared with the PS19 group (Figure 6). Notably, hyperphosphorylated human tau was predominantly localized within the insoluble protein fraction of the brain, indicating its incorporation into aggregates. They also displayed more Ub conjugates, indicating proteasomal insufficiency (Figure 7). Since the genetic blockade of p-S14-Rpn6 was the only difference between the two groups, the p-S14-Rpn6 was responsible for the lesser tau accumulation and therefore preserved proteostasis observed in PS19 mice compared with the PS19::S14A/A mice. However, S14-phopshorylated Rpn6 was not increased in PS19 mice compared to WT mice, which is in agreement with our previous findings from a model of cardiac proteinopathy, where overexpression of a disease-linked misfolded protein in cardiomyocytes does not increase S14-phosphorylated Rpn6 and in fact decreases it (22). Nevertheless, these findings implicate that there is likely a room for p-S14-RPN6 augmentation to prevent or reduce AD tauopathy.

While the role of cAMP/PKA augmentation was not directly explored in the current study, previous research has shown that cAMP/PKA is responsible for phosphorylating S14-Rpn6 (22). Phosphodiesterase (PDE) inhibitors are a type of drug that prevents phosphodiesterases from degrading cAMP and/or cGMP and these have been investigated as treatments for AD. A clinical study examined the effect of Cilostazol, a PDE3 inhibitor, as an add-on medication in AD patients and found reduced odds of decline in cognitive function (47). Another more relevant study, administered rolipram, a phosphodiesterase 4 inhibitor, in AD model mice and observed increased proteasome activity, which reduced tauopathy, and consequently improved cognitive function (16). The study speculated that this could be via p-S14-Rpn6 but did not directly test this hypothesis. Additionally, this study only observed drug treatment effects on the brain, leaving how treatment would affect tauopathy of the heart untested. Hence, future research is necessary to investigate whether augmentation of p-S14-Rpn6, either through genetic or pharmacological strategies, can attenuate tau pathology and improve neurodegenerative outcomes in both PS19 and other AD mouse models.

In conclusion, our study has provided compelling evidence that phosphorylation of S14-Rpn6 has a critical role in preserving proteasomal function and maintaining proteostasis in tauopathy-based AD. Our study also underscores the importance of p-S14-Rpn6 of the 26S proteasome as a potential therapeutic target for treating both the brain and heart pathology of AD.

## METHODS

### Sex as a biological Variable

Our study examined both male and female animals in an approximately 1:1 ratio.

### Animals

The creation and validation of the Rpn6^S14A^ knock-in (S14A) mouse model were described previously (22). In the S14A mice, the serine14 codon of *Rpn6* is mutated to an alanine to block the normal phosphorylation of serine 14. The S14A mouse model was crossbred with the Tau P301S (PS19) mouse [B6;C3-Tg(Prnp-MAPT*P301S)PS19Vle/J; Jackson Laboratories strain #008169; RRID: IMSR_JAX:008169]. PS19 mice overexpress the T34 isoform of microtubule-associated protein tau with one N-terminal insert and 4 microtubule-binding repeats encoding the human P301S mutation, all driven by the mouse prion protein promoter (24). The animals were given ad-lib access to food and water and housed in specific pathogen free control rooms with optimal temperature (22-24°C) and 12 hr light/dark cycle. The protocols for animal care and use in this study have been approved by the University of South Dakota Institutional Animal Care and Use Committee.

### Open Field Test

In a dimly lit room, mice were placed in the center of an open field arena (52×33.5×30 cm) and allowed to freely explore for 10 min. The Ethovision XT software (Noldus) tracked the distance travelled and velocity for each mouse. The arena was cleaned between mice (48).

### Novel Object Recognition Test

On the first day of the object recognition test, mice were placed into the empty test box (43×30×38 cm) for 10 min to habituate to testing conditions. On the second day, two identical objects were placed in the test box and the mice explored for 10 min. After 5 minutes of rest in their original cage, one of the objects was replaced with a new similar sized object and the mice again explored the box for 3 minutes. The movement of the mice was tracked with the Ethovision XT software (Noldus). Exploration was defined as time spent near an object and the number of approaches to an object. A mouse’s recognition of the new object was determined by the recognition index (RI) (RI = T_novel_ / T_novel_ + T_familiar_) and the discrimination index (DI) (DI = (T_novel_ - T_familiar_)/(T_novel_ + T_familiar_)). Total distance was also recorded. The arena was cleaned between mice (48).

### Rotarod Test

The Rotarod apparatus (Stoelting) consisted of an accelerating rod (4 rpm–12 rpm) separated into five lanes. Animals were tested five at a time with each animal placed on a random lane. The mice were trained for three successive days with 3 trials each day. The maximum trial length was 700 sec. The fourth day was testing day, with 3 trials for each mouse. Time on the rod was recorded and the average of three trials was used in the analysis. The rod, lanes, and chamber were cleaned between mice (48) .

### Echocardiography

Echocardiography was performed at 3, 6, and 9 months of age. Mice were anesthetized with isoflurane at 4% initially then 1.5% for maintenance. A transthoracic short-axis M-mode echocardiograph (22), pulse wave, and tissue doppler velocities (49) were recorded with the 40-MHz probe of the VisualSonics Vevo 3100 imaging system (FUJIFILM Sonosite). Heart rate (HR), left ventricular mass (LVmass), left ventricular anterior wall thickness (LVAW), left ventricular posterior wall thickness (LVPW), left ventricular end-diastolic volume (LVEDV), left ventricular end-diastolic volume (LVESV), left ventricular internal diameter at end diastole (LVID;d), left ventricular internal diameter at end systole (LVID;s), ejection fraction (EF), fractional shortening (FS), stroke volume (SV) and cardiac output (CO), E/e’, E/A and deceleration time were determined using the Vevo Lab software (FUJIFILM Sonosite).

### Isolation of Hippocampus

To isolate the hippocampus from mice for downstream western blot analyses, we used a protocol published by Jaszcyk *et al*. (50), with some minor modifications. This method enabled us to dissect the hippocampus tissue out of mouse brain after removing adjacent tissue contamination. The technique helped us avoid tissue distortions, allowing us to obtain pure hippocampal tissue suitable for downstream molecular experiments performed on the entire hippocampus.

### Total Protein Extraction and Western Blot Analysis

Total protein extraction and western blot analysis were performed according to previously described methods (22). Briefly, heart ventricular tissue and hippocampal tissue from 9 month old mice were snap-frozen in liquid nitrogen and stored at -80 °C. Ventricular tissue was homogenized in lysis buffer (pH 6.8, 50mM Tris-HCl, 2% SDS, 10% glycerol, protease, and phosphatase inhibitor cocktail, [#HB9105, HelloBio]) with the Bead Blender (Next Advance) then boiled for 5 min and centrifuged at 10,621x g and 4°C for 20 min. Hippocampal tissue was homogenized in RIPA buffer (pH 7.6, 25 mM Tris, 150 mM sodium chloride, 1% NP-40, 1% sodium deoxycholate, 0.1% SDS, #786-489 G-Biosciences).

Supernatant protein concentration was determined with the bicinchoninic acid assay (#23222, #23224, Thermo Fisher Scientific). For each mouse, 20 μg of protein was fractionated in 10% stain-free SDS-PAGE gels and transferred to PVDF membranes (#1620177, Bio-Rad Laboratories) with the Mini Trans-Blot Cell (Bio-Rad Laboratories) for all blots, except pS14-Rpn6, which required 60 μg. All membranes were blocked with 2.5% bovine serum albumin (#A30075, Research Products International) for 1 hr at room temperature, except pS14-Rpn6 which was blocked with ECL advanced blocking agents (#RPN418, Cytiva). Membranes were then incubated with primary antibody (Neu N #A19086, GFAP #A14673, IBA1 #A19776, pTauS202/T205 #AP0894, Abclonal; A10 #sc-390476, PHF 13 #sc-92275, Synaptotagmin #sc-393392, Santa Cruz; PSD-95 #2507S, Ubiquitin (P37) #58395, Cell Signaling; Rpn6 #MBS9605099, MyBioSource; Ubiquitin Lys48-Specific #05-1307, Millipore Sigma; and CP27 and pS14-Rpn6 were custom made and generously donated) at 4 °C overnight and then incubated in horseradish peroxidase-conjugated secondary antibody (#115035003, #111035003, Jackson ImmunoResearch) at room temperature for 1 hr. Blot images were taken with the ChemiDoc MP (Bio-Rad Laboratories) imaging system after incubation with chemiluminescent substrate (#34578, Thermo Fisher Scientific). Blots were normalized against total in-lane protein with the ImageLab software (Bio-Rad Laboratories) as previously described (51).

### Extraction of NP-40-soluble and NP-insoluble Fractions

The protocol for the fractionation of NP-40 soluble and insoluble proteins was described previously (22). In brief, frozen ventricular myocardium and hippocampus were homogenized in lysis buffer and incubated at 4 °C for 30 min and then centrifuged for 15 min at 17,000 g and 4 °C. The supernatant (NP-40 soluble fraction) was collected and added to SDS boiling buffer (pH 8.0, 6% SDS, 20 mM Tris-HCl, 150 mM DTT) and boiled for 5 min. The pellet (NP-40 insoluble fraction) was washed in PBS and put in cell pellet buffer (pH 8.0, 20 mM Tris-HCl, 15 mM MgCl2, 2 mM DTT, protease and phosphatase inhibitor cocktail, [#HB9105, HelloBio]) and incubated for 30 min at 4 °C. The insoluble fraction was put into SDS boiling buffer and boiled for 5 min. Protein concentrations were determined with a reducing agent-compatible BCA kit (#23252, Thermo Fisher Scientific).

### RNA isolation and qPCR

The apex of each ventricular myocardium was stored in RNALater (#AM7024, Thermo Fisher Scientific). The tissue was homogenized with the Bead Blender (Next Advance) and mRNA was extracted with TRI Reagent (TR 118, Molecular Research Center) mRNA concentrations were determined with the NanoDrop 2000 UV Spectrophotometer (Thermo Fisher Scientific) as previously described (51). The High-Capacity cDNA Reverse Transcription Kit (#4368814, Applied Biosystems) was used to generate cDNA by following the manufacturer’s protocol. For the qPCR assay, each well contained 2 μL of adjusted cDNA sample solution, 0.2 μM of both forward and reverse primers, 10 μL of 2X SYBR Green master mix (#RK21203, Abclonal), and a nuclease free water to 20 μL total volume. The samples were measured, in triplicate, in a 96 well plate (#N8010560 Thermo Fisher Scientific). The reactions were measured in the QuantStudio 6 PCR System (Thermo Fisher Scientific) with the program: 50 °C for 2 min, 95 °C for 10 min, followed by 40 cycles of 95 °C for 15 sec and 60 °C for 60 sec. Target gene expressions were normalized to *Gapdh* using the 2-ΔΔCt method. The genes investigated and their primers are *Collagen 1* (Forward 5’-GCCACTGCCCTCCTGACG-3’; Reverse 5’-GCCATCTCGTCCAGGGG-3’), *Nppa* (Forward 5’-GGAGGAGAAGATGCCGGTAGA-3’; Reverse5’-GCTTCCTCAGTCTGCTCACTCA-3’), *Nppb* (Forward 5’-CTGCTGGAGCTGATAAGAGA-3’; Reverse 5’-TGCCCAAAGCAGCTTGAGAT-3’), and mouse *Gapdh* (Forward 5’-ATGACATCAAGAAGGTGGTG-3’; Reverse 5’-CATACCAGGAAATGAGCTTG-3’).

### Fluorescence staining and confocal microscopy

We followed the previously reported methods for Immunofluorescence staining and confocal microscopy (22). In brief, brain and ventricular myocardium were fixed in 4% paraformaldehyde, embedded in Tissue-Tek O.C.T. Compound (Sakura), and then cryosectioned into 5 μm sections with the CM 1590 cryostat (Leica Microsystems). The sections were blocked in 5% BSA for 1 hour and then incubated with primary antibody (GFAP #A14673, IBA1 #A19776, Abclonal; Synaptotagmin #sc-393392, Santa Cruz; PSD-95 #2507S, Cell Signaling) overnight at 4 °C and then incubated with a fluorescent secondary antibody (#A12380, Thermo Fisher Scientific; #20478, Cayman Chemical) for 2 hr. DAPI (#0100-20, Southern Biotech) was used to stain nuclei. The sections were imaged with the TCS SP8 confocal microscope (Leica Microsystems) and analyzed with ImageJ software (NIH).

### Picrosirius red staining

Before tissue collection, mice were perfused with PBS and then 10% formalin PBS under isoflurane anesthesia. The hearts were incubated in 10% formalin PBS at 4 °C for 24 h.

The hearts were then dehydrated and embedded into paraffin blocks. The blocks were cut into 5 μm sections with an RM 2155 microtome (Leica) and set on charged slides. Slides were deparaffinized and rehydrated then fixed in Bouin’s solution, stained in Wiegert’s iron hematoxylin and picrosirius red stain solutions, and mounted (52). Slides were imaged with the Olympus IX71 microscope (Olympus) and analyzed with ImageJ software (NIH).

### Statistical Methods

GraphPad Prism version 9.0 (GraphPad Software) was used for all graphs and statistical tests. Data are represented by mean±SEM. Statistical methods for each experiment are specified in each figure legend. In summary, normally distributed data were analyzed with one-way ANOVA followed by Tukey’s test for multiple comparisons except in Figures 3B and 3F, 4B∼4E, 6B∼6E, and 7B where technical replicates are included and accordingly a nested one-way ANOVA was utilized. Non-parametric data were analyzed with Kruskal Wallis followed by Dunn’s test.

## Supporting information

Supplemental Figures S1 through S4

## Data availability

Values for all data points in the graphs are reported in the supporting data value.xls file. Other data from this study are available upon reasonable request.

## Funding

This work is supported in part by NIH grants P20RR17662-019003, RF1AG072510, R01HL072166, and R01HL153614.

## Notes

### Competing Interest Statement

The authors have declared no competing interest.

