## Supplemental Figures S1 through S4 for "Ser14-phosphorylated Rpn6 Limits Proteostasis Impairment and Pathology in Both Brain and Heart of Tauopathy Mice"

### Supplementary Figures 1-4

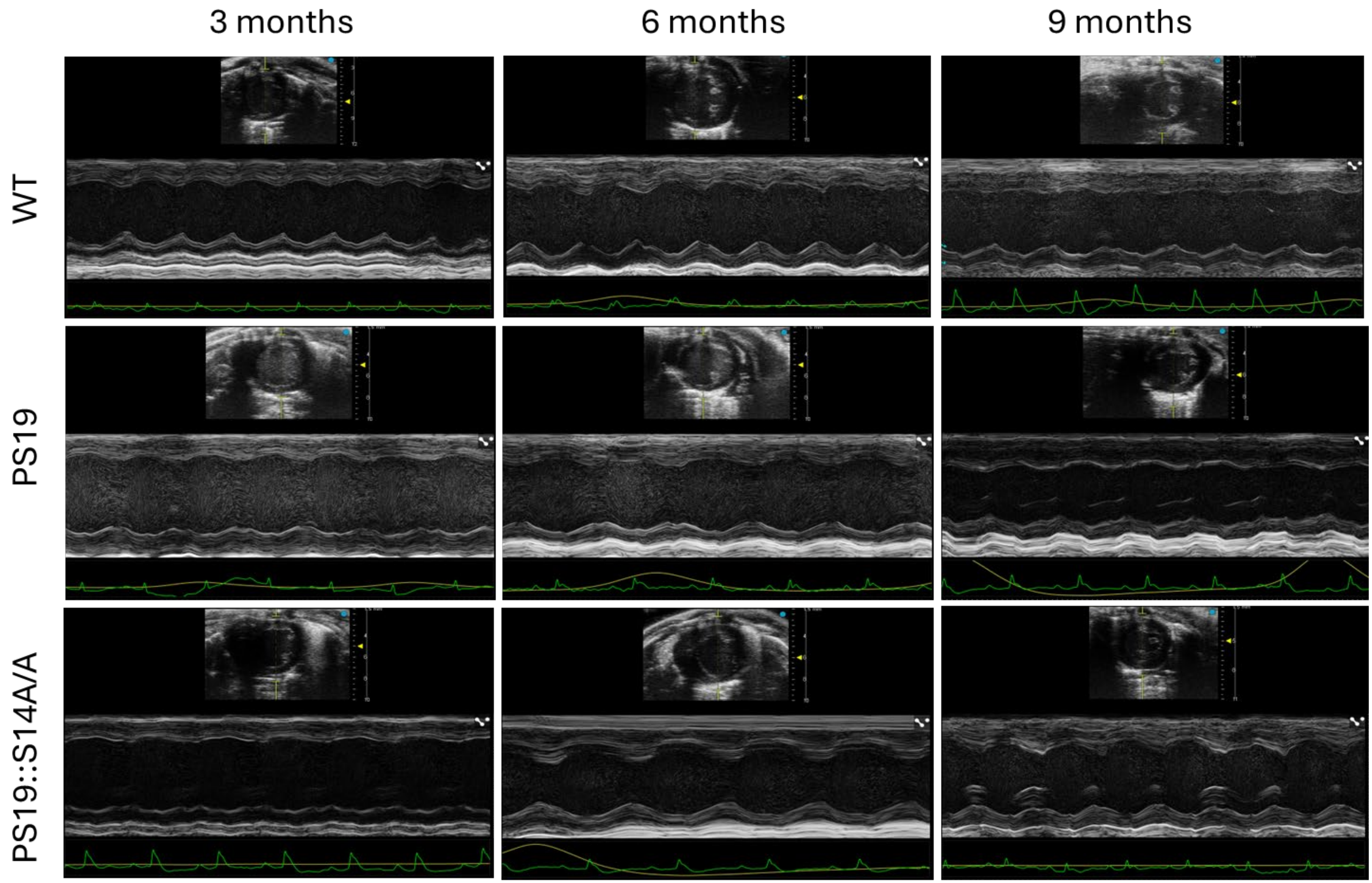


**Figure S1. Representative images of echocardiography**. Mice of indicated genotypes underwent echocardiography at 3, 6, and 9 months.

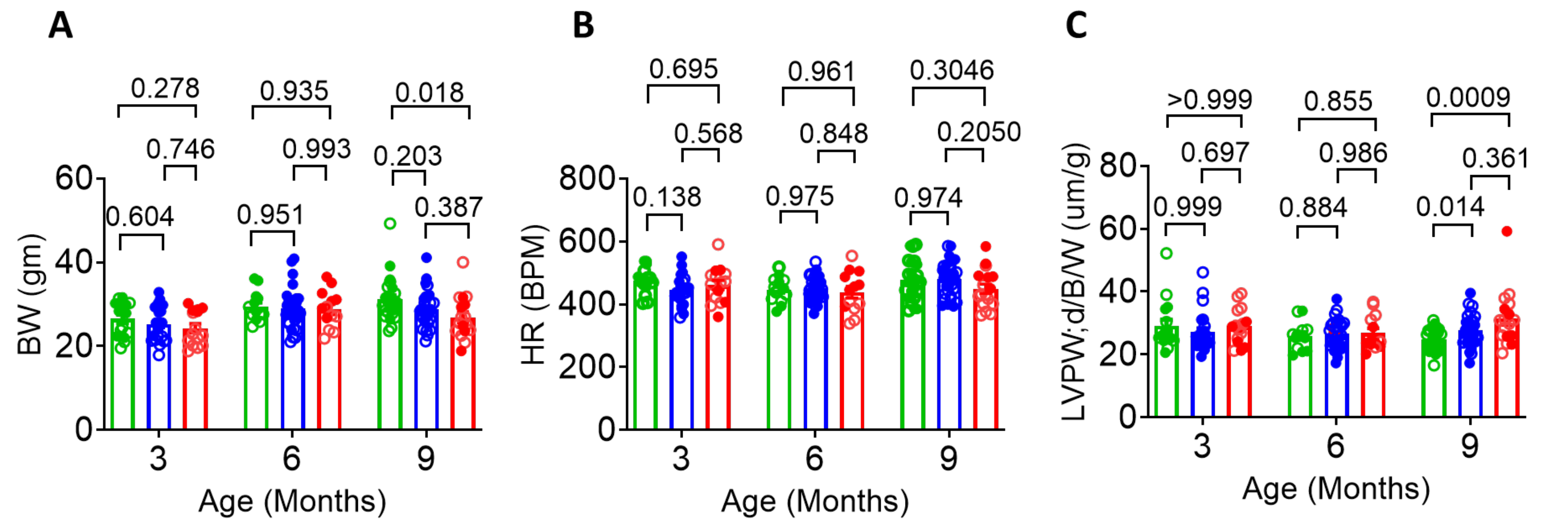


**Figure S2. Additional parameters of M-mode echocardiography**. Mice of indicated genotypes underwent echocardiography at 3, 6, and 9 months of age as described in Figure 2 of main text.

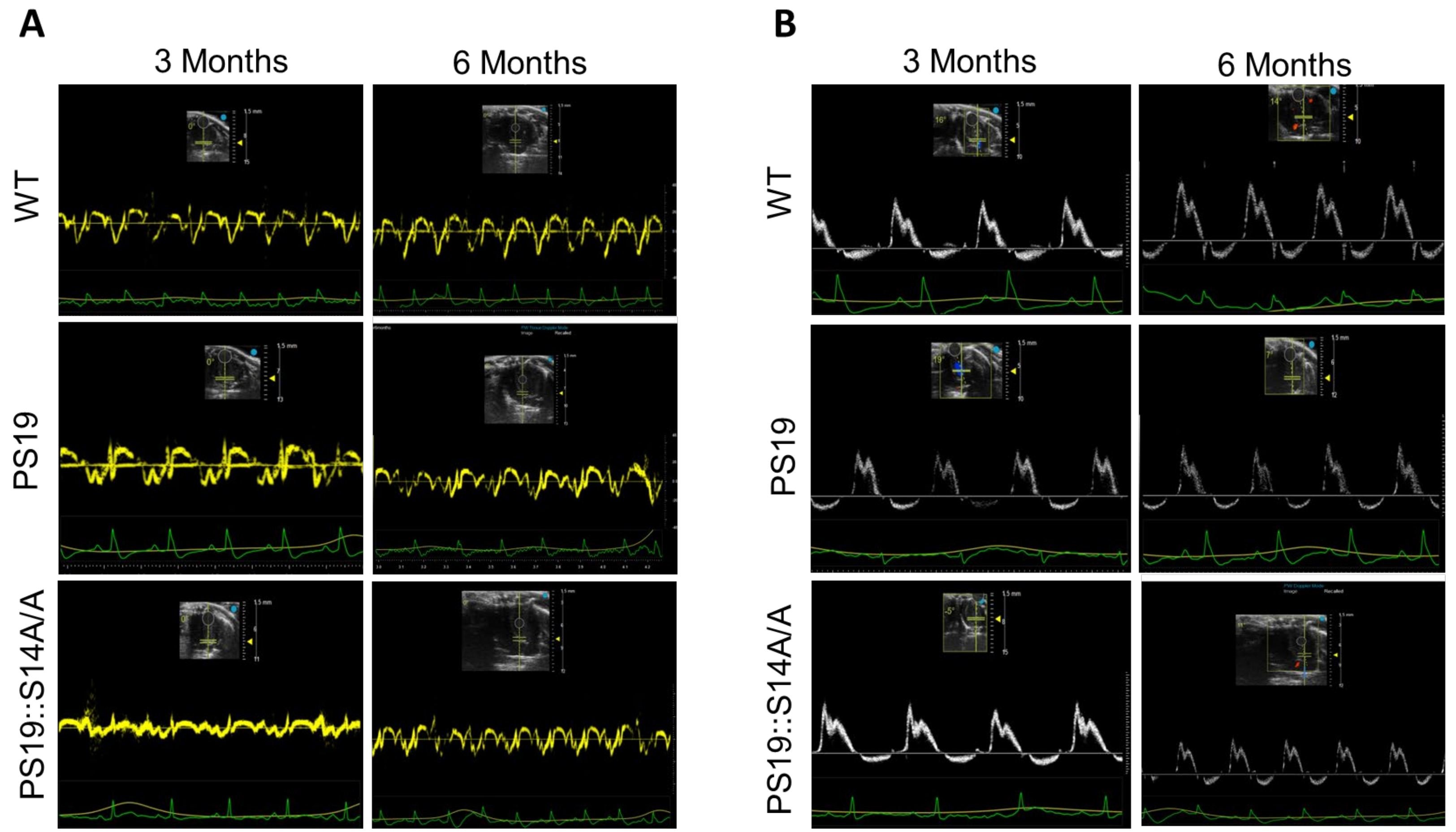


**Figure S3**. Representative echocardiographic images of flow tissue Doppler at the level of the mitral valve annulus (**A**) and pulsed-wave Doppler of the mitral valve (**B**).

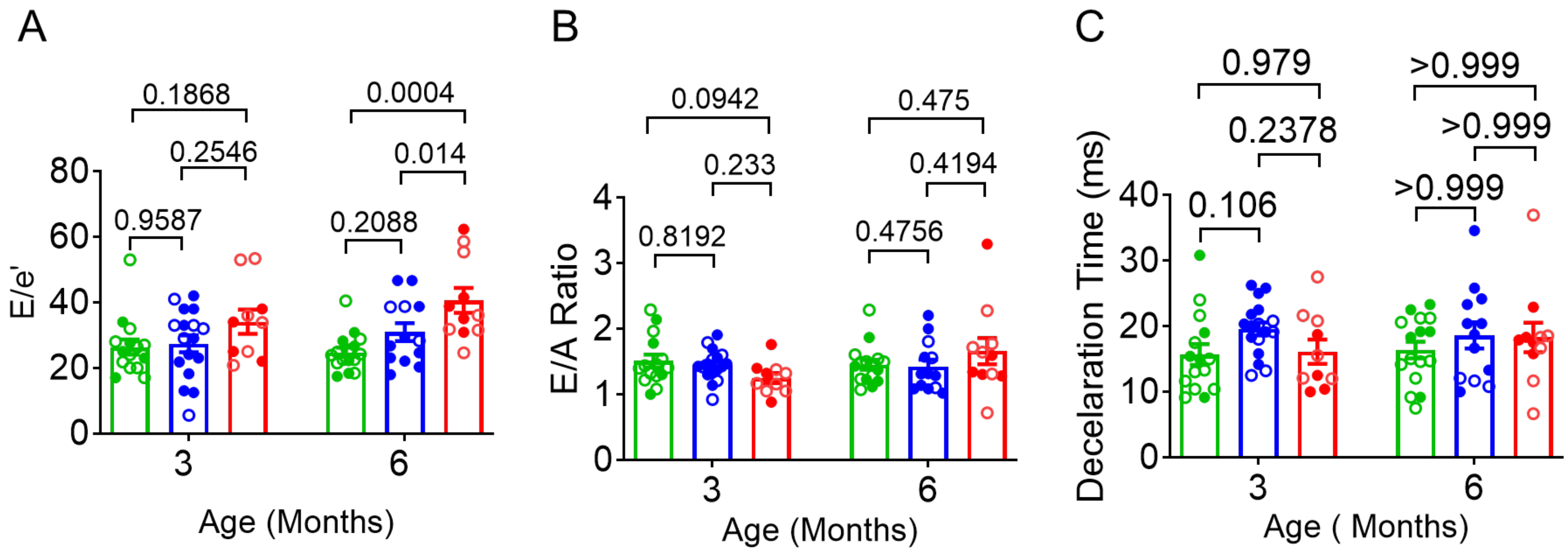


**Figure S4**. Diastolic function assessment from flow tissue Doppler at the level of the mitral valve annulus and pulsed-wave Doppler of the mitral valve. (**A**) E/e' ratio, here the early diastolic peak velocity of blood flow across the mitral valve (E) was measured using pulsed-wave Doppler and the early diastolic peak velocity of the mitral annulus (e') was measured using tissue Doppler imaging. **(B**) E/A ratio, and (**C**) Deceleration Time for 3 and 6-month-old mice. WT (3m: males=5, females=10; 6m: males=6, females=10); PS19 (3m: males=11, females=6; 6m: males=10, females=3; PS19::S14A/A (3m: males=4, females=6; 6m: males=5, females=6) for diastolic function.
